# Multi_CycGT: a DL-based multimodal model for membrane permeability prediction of cyclic peptides

**DOI:** 10.1101/2023.06.20.545822

**Authors:** Lujing Cao, Zhenyu Xu, Tianfeng Shang, Chengyun Zhang, Xinyi Wu, Yejian Wu, Silong Zhai, Liefeng Ma, Hongliang Duan

**Affiliations:** College of Pharmaceutical Sciences, Zhejiang University of Technology, Hangzhou, 310014, P. R. of China; AI department, Hangzhou Highline Therapeutics. Inc, Hangzhou 310014, China; State Key Laboratory of Drug Research, Shanghai Institute of Materia Medical (SIMM), Chinese Academy of Sciences, Shanghai, 201203, P. R. of China

## Abstract

As a highly versatile therapeutic modality, cyclic peptides have gained significant attention due to their exceptional binding affinity, minimal toxicity and capacity to target the surface of conventionally “undruggable” proteins. However, the development of cyclic peptides with therapeutic effects by targeting intracellular biological targets has been hindered by the issue of limited membrane permeability. In this paper, we have conducted an extensive benchmarking analysis of a proprietary dataset consisting of 6941 cyclic peptides, employing machine learning and deep learning models. In addition, we propose an innovative multimodal model called Multi_CycGT which combines a Graph Convolutional Network (GCN) and a Transformer to extract 1D and 2D features. These encoded features are then fused for the prediction of cyclic peptide permeability. The cross-validation experiments demonstrate that the proposed Multi_CycGT model achieved the highest level of accuracy on the test set, with an accuracy value of 0.8206 and an AUC value of 0.8650. This paper introduces a pioneering deep learning-based approach that demonstrates enhanced effectiveness in predicting the membrane permeability of cyclic peptides. It also represents the first attempt in this field. We hope that this work will help to accelerate the design of cyclic peptide active drugs in medicinal chemistry and chemical biology applications.

## Introduction

The last decade has seen an increase in the number of peptide drugs approved by regulatory authorities and cyclic peptide becomes a potential candidate gaining much attentions^1^. The formation of cyclic peptides observed in nature often occurs through reactions involving two reactive groups. These reactions include thiolactonization, lactamization, lactonization, or disulfide bridge formation^2^. Cyclic peptides also can be divided into different classes: head-to-tail, head-to-side and side-to-side. Such ring structure has several advantages over traditional linear peptides: (1) the formation of hydrogen bonds within the ring peptide molecule greatly reduces the molecular polarity, which makes its metabolic stability and bioavailability much higher than that of straight-chain peptides; (2) ring peptides have a clear degree of structural pre-organization, which can better fit the receptor; (3) ring peptides generally have a large surface area, which makes them highly affinity and recognition specific^3-7^. On average, the FDA has approved one macrocyclic peptide drug per year over the past decade^8^. However, it is worth noting that among these approved macrocyclic peptide drugs, only romidepsin^9^ that utilized for the treatment of lymphoma, specifically acts on an intra-cellular target. This highlights the limitation of macrocyclic peptides in penetrating cell membranes and interacting with intracellular targets, which has been a blocking stone in this field.

Currently, the strategies such as N-methylation^10, 11^, amide-to-ester substitution^12^, side-chain modification^13, 14^, and D-amino acid substituents^15^ could increase the membrane permeability of cyclic peptides. However, it still takes a lot of time and money for researchers to test the membrane permeability of modified cyclic peptides. In recent years, the membrane permeability of cyclic peptides has been predicted by a number of statistical or machine learning methods^13, 16-20^. Rezai *et al*. used a continuous solvation model to predict the passive membrane permeability of 11 cyclic peptides with an *R*^2^ of 0.96. The model quantitatively estimated the solvation free energy loss which was correlated with PAMPA data. The study by Wang^21^ discussed the relationship between several physical parameters and membrane permeability and finally developed a predictive model. This method described the correlation between hydrogen bonding potential characteristics and membrane permeability, and was validated by molecular dynamics-based simulations.

Compared to traditional statistical and machine learning methods, deep learning (DL)models can automatically extract more complex and effective potential features from datasets, which may facilitate new insights into cyclic peptide membrane permeability. Therefore, this article is contributed to predict membrane permeability effectively by utilizing DL-based models. In fact, building DL-based predictive models for such task has been a serious challenge, due to the limited and heterogeneous corresponding experimental data. Thanks to the appearance of the CycPeptMPDB^22^, making computational membrane permeability prediction comes true. The creators of this database had collected 7334 structurally distinct cyclic peptides, and its corresponding membrane permeability information from 45 published papers and 2 patents of pharmaceutical companies. In addition to the experimental data, the database system contains supporting information such as SMILES representations of cyclic peptides, amino acid representations, allowing users to analyze and utilize it in multiple dimensions.

In this work, we used this database to develop a model termed Multi_CycGT (Multimodal model combined with GCN and Transformer for membrane permeability prediction of cyclic peptide) to reliably predict the membrane permeability of cyclic peptides. This model is equipped with two main architectures: Transformer and GCN. The GCN module uses the cyclic peptide atomic-level properties and interatomic connectivity information (2D feature) to dynamically learn and translate them into appropriate vectors. The Transformer modules encodes the SMILES sequence features (2D feature) of cyclic peptides through a text-based translation task. Finally, these different morpho-logical features are concated with the physicochemical property features through the fully connected (FC) layer for cyclic peptide membrane permeability classification task (Figure 1). The results show that the Multi_CycGT model significantly outperforms other machine learning or deep learning approaches in five metrics: accuracy (ACC), precision (Pre), F1-score (F1), recall score (Recall) and area under the curve (AUC). In addition, we performed ablation experiments and feature analysis. The main contribution of our study is revealing a fact that characterization methods integrating different dimensional features can improve the accuracy of prediction of cyclic peptide permeability. To our best knowledge, this article is the first work on DL-based predicting the membrane permeability of cyclic peptides. We hope that our work can contribute to the progress of deep learning in cyclic peptide drug design and discovery, and the understanding of the descriptors or features related to cyclic peptide transmembrane properties to inform future work on designing rational transmembrane cyclic peptide drugs.

**Figure 1.**
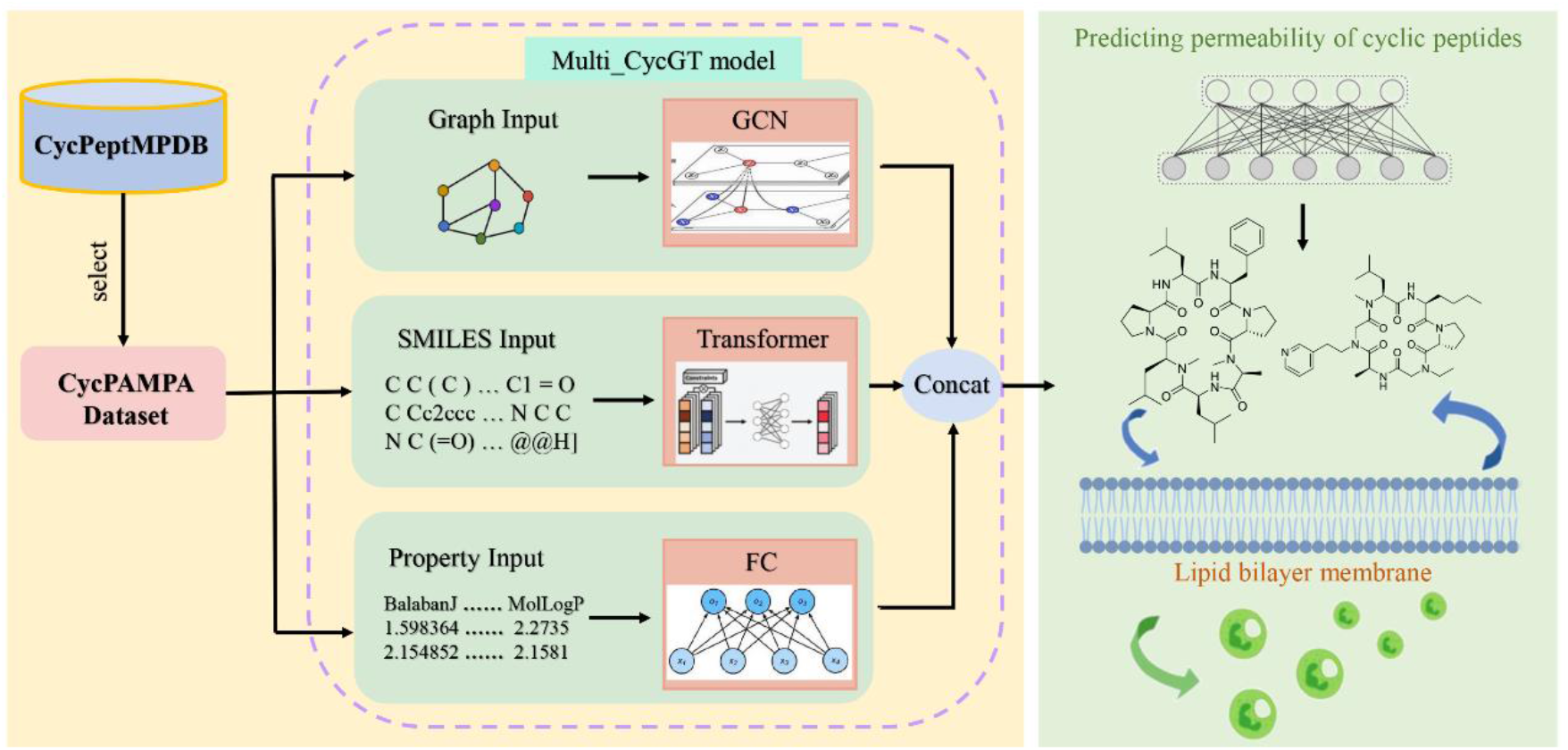
An overview of predicting membrane permeability of cyclic peptides. The inputs of this model involve three categories of features: 1D SMILES and property descripors, and 2D Graph. The GCN absorbs the graph information of cyclic peptide. The Transformer module is trained on SMILES. A Fully connected (FC) layer focus on properties such as MolLogP, BalabanJ. Those different features are concated and then delivered to the final classifier.

## Results and discussion

Before displaying the performances of models in permeability prediction, we analyzed the dataset distribution of properties, which may help readers getting further understanding of our study. In this study, we utilized the CycpepMPDB_PAMPA dataset, which comprises a collection of 6941 cyclic peptides sourced from the literature. These peptides exhibit a range of sequence lengths, spanning from 2 to 15 amino acids. It is noteworthy that more than 90% of the cyclic peptides are comprised of non-natural amino acids. This indicates that these peptides are chemically modified or designed to incorporate non-natural building blocks, possibly to enhance their stability, binding affinity, or other desired properties. However, it is important to acknowledge that the conformation and membrane permeability mechanisms of these non-natural cyclic peptides have not been extensively studied or clearly elucidated. The limited knowledge in this regard highlights the need for further investigation and understanding of the structural and functional characteristics of these modified cyclic peptides. As shown in Figure 2, these peptides are generally much larger than traditional small molecule drugs, and most have high molecular weight (only 1.4% molecular weight <500 Da). In terms of membrane permeability of cyclic peptides, tPSA which may reflect the transport properties of drugs in the cell, also shows no strong correlation. In other respects, they largely conform to Lipinski’s five rules (Ro5)^23^. In conclusion, unlike traditional medicinal chemistry compounds, it is necessary to explore the membrane permeability of cyclic peptides and its relationship with property features outside the Ro5 space using the DL-based approaches.

**Figure2.**
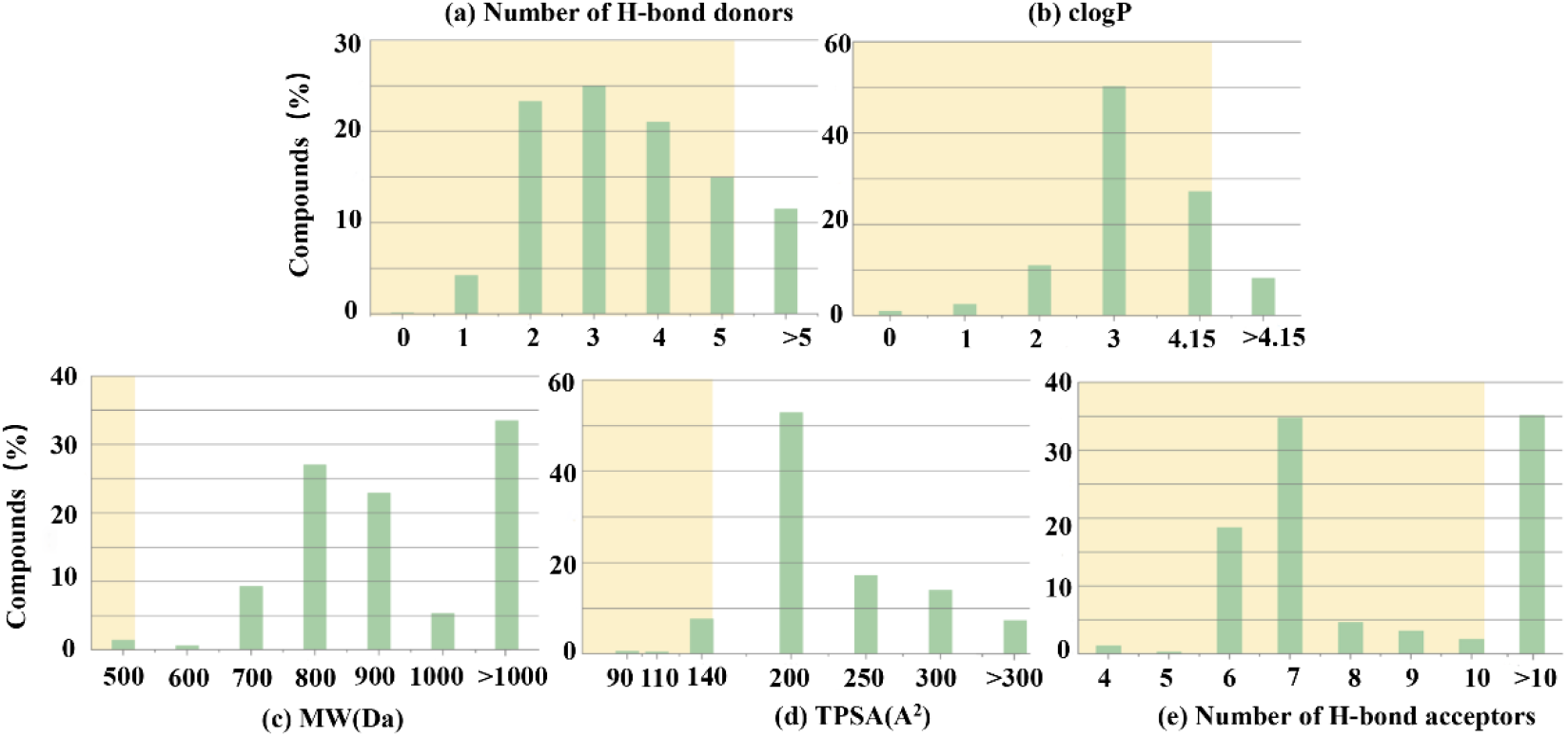
The histogram shows the property distributions of 6941 cyclic. The highlighted areas in blue show compounds that are compliant with the Lipinski for the property in question. Properties are as follows: number of H-bond donors, TPSA, MW, number of H-bond acceptors and clogP.

### Model performance comparison

We implemented several models involving different mechanisms to predict membrane permeability of cyclic peptides (detailed information about models can be founded in Method). The 10-fold cross-validation results (Table 1) show that the SVM, KNN, CNN, Transformer, LSTM, GCN, and Multi_CycGT models have an accuracy value of 0.7994, 0.7521, 0.7848, 0.7532, 0.7141, 0.7777, and 0.8206, respectively. In this experiment, the Multi_CycGT model performs best, followed by the SVM-based classifier, then the CNN. Moreover, the Multi_CycGT model achieves the highest F1-score and precision among the evaluated models, indicating its superior ability to balance both sensitivity and positive predictive value in predicting cyclic peptide membrane permeability. These findings highlight the effectiveness of the Multi_CycGT model in accurately predicting and assessing the membrane permeability of cyclic peptides. Contrary to expectations, the SVM have a best recall performance. Although the accuracy rate can determine the overall prediction correctness, the accuracy rate alone may cause some one-sided error when the positive and negative samples of the dataset are unbalanced. To avoid such case, we adopt the AUC as a main metric in our work. Inspiringly, Figure 3 displays that the Multi_CycGT model has an AUC value of 0.8650, which is higher than any other methods. This suggests that the Multi_CycGT model is adept at capturing and distinguishing the positive instances, even in the presence of such bias. Overall, the Multi_CycGT model displays a powerful ability to predict the membrane permeability of cyclic peptides on the CycpepMPDB_PAMPA dataset. It consistently outperforms almost all other methods in terms of most of evaluation metrics.

**Table 1.** The performances of various models with different metrics on the test set. These results are all average values of 10-fold cross-validation.

| Model | ACC | Pre | F1_Score | Recall |
| --- | --- | --- | --- | --- |
| SVM | 0.7994 | 0.8223 | 0.8606 | **0.9032** |
| KNN | 0.7521 | 0.7744 | 0.8331 | 0.9018 |
| CNN | 0.7848 | 0.8126 | 0.8493 | 0.8909 |
| Multi_CycGT | **0.8206** | **0.8600** | **0.8708** | 0.8820 |
| Transformer | 0.7532 | 0.7710 | 0.8395 | 0.9234 |
| LSTM | 0.7141 | 0.8200 | 0.8095 | 0.8980 |
| GCN | 0.7777 | 0.8239 | 0.8158 | 0.8752 |

**Figure 3.**
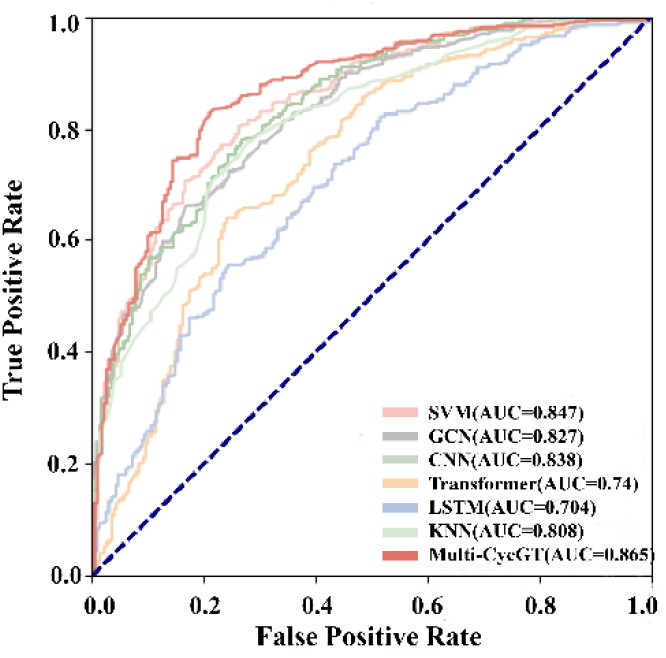
The AUC performance of machine learning and deep learning on the test set Evaluation of Multi_CycGT model capability to predict cyclic peptides’s membrane permeability.

### Ablation experiment

In this study, the physicochemical properties of the FC layer, Transformer text features and GCN features are concated to predict membrane permeability of cyclic peptides. Here, we conducted an ablation study to explore the effect of these different modules. Under the same experimental setup, we adopt two simplified variants of Multi_CycGT model for predicting cyclic peptide’s membrane permeability:

1. Trans_FC: by only containing the transformer architecture and a FC layer.
2. GCN_FC: by only containing the GCN architecture and a FC layer.

Figure 4 shows the performance comparison of ACC, Pre, F1, Recall, and AUC for the Multi_CycGT model and different variants on the test set. It clearly demonstrates that the feature extraction modules of both Trans_FC and GCN_FC have a significant impact on the permeability prediction ability of models. In 10-fold experiments, the average ACC of Trans_FC, GCN_FC and Multi_CycGT reach 0.7917, 0.7889 and 0.8206 respectively. These two simplified variants of Multi_CycGT models gain comparable accuracy in our prediction task. The similarity of these results between Trans_FC and GCN_FC models means that the 1D feature (SMILES text) is as same important as 2D feature (cyclic peptide graph) in this task. The observed trend in both the Pre and F1 metrics remains consistent. Notably, Trans_FC 、 GCN_FC and Multi_CycGT have average Recall values of 0.9049, 0.8791, and 0.8820, respectively. However, the Trans_FC model demonstrates a notably stronger performance in predicting positive samples. By combining these two innovative elements, we notice that our proposed multimodal model can better fuse the characteristic properties of cyclic peptides in terms of permeability, and the permeability prediction of cyclic peptides is obviously better compared to the Trans_FC and GCN_FC. This phenomenon also reveals that synergy between 1D and 2D features. From the biological point of view, the adoption of some strategies may be crucial to improve the pharmacokinetic properties of cyclic peptide drugs, and the information of these local structures can be captured by our multimodal model.

**Figure 4.**
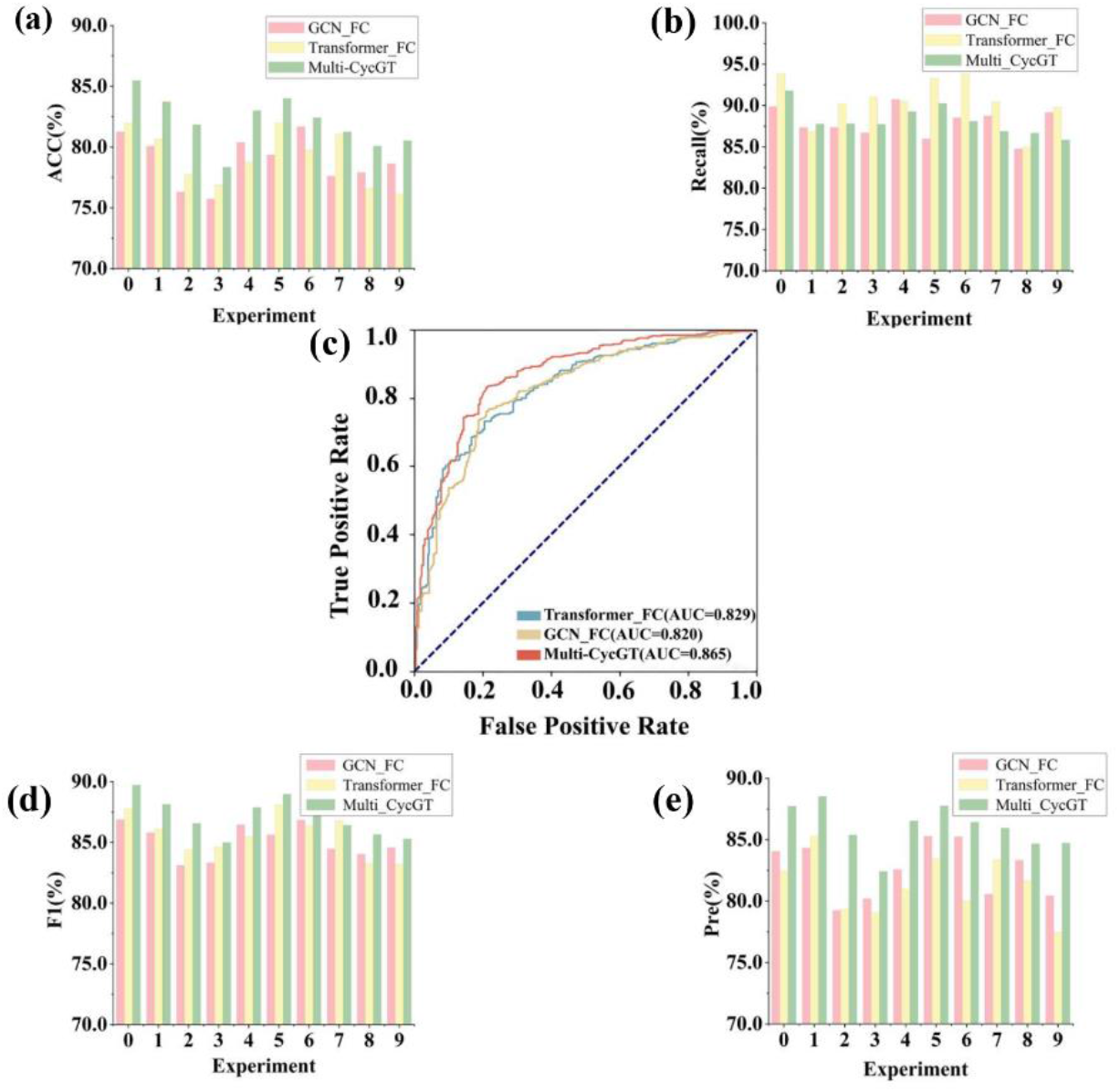
The performances of variants of Multi_CycGT models in 10-fold cross-validation with different metrics.

### Model Performance with different peptide lengths

The dataset contains cyclic peptide sequences ranging in length from 2 to 15, but the majority of sequence lengths are 6 and 7. This is probably due to the fact that shorter cyclic peptides prefer to be a more compact spatial conformation in response to the membrane permeability property. The distribution of different peptide lengths in the dataset is shown in Figure 5. Faced with such data contribution, we wonder if the peptides sequences may affect the prediction ability of our model. In order to address this problem, we discuss results of models with different sequence lengths in this section.

**Figure 5.**
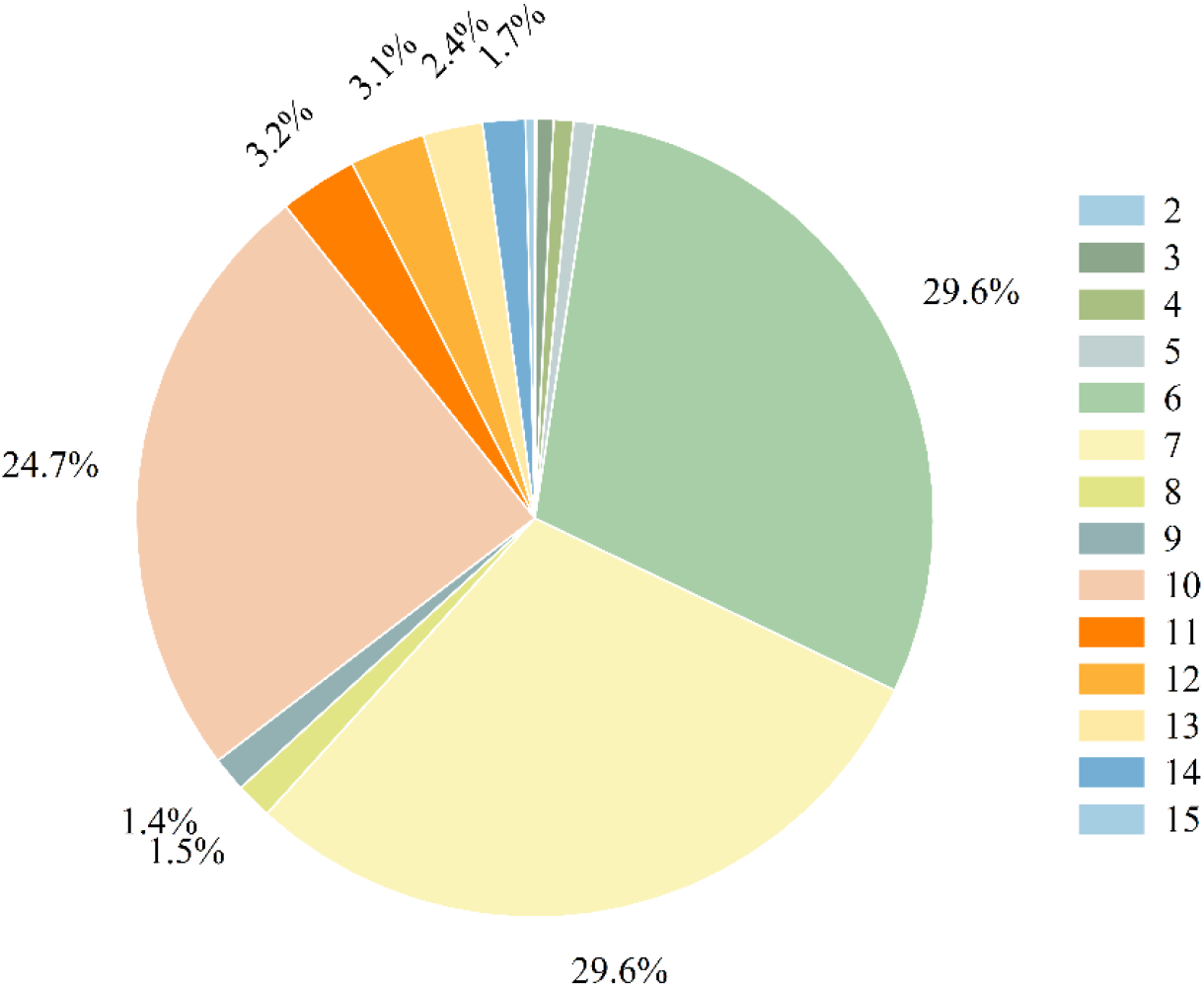
The distribution ratio of cyclic peptide lengths.

**Figure 6.**
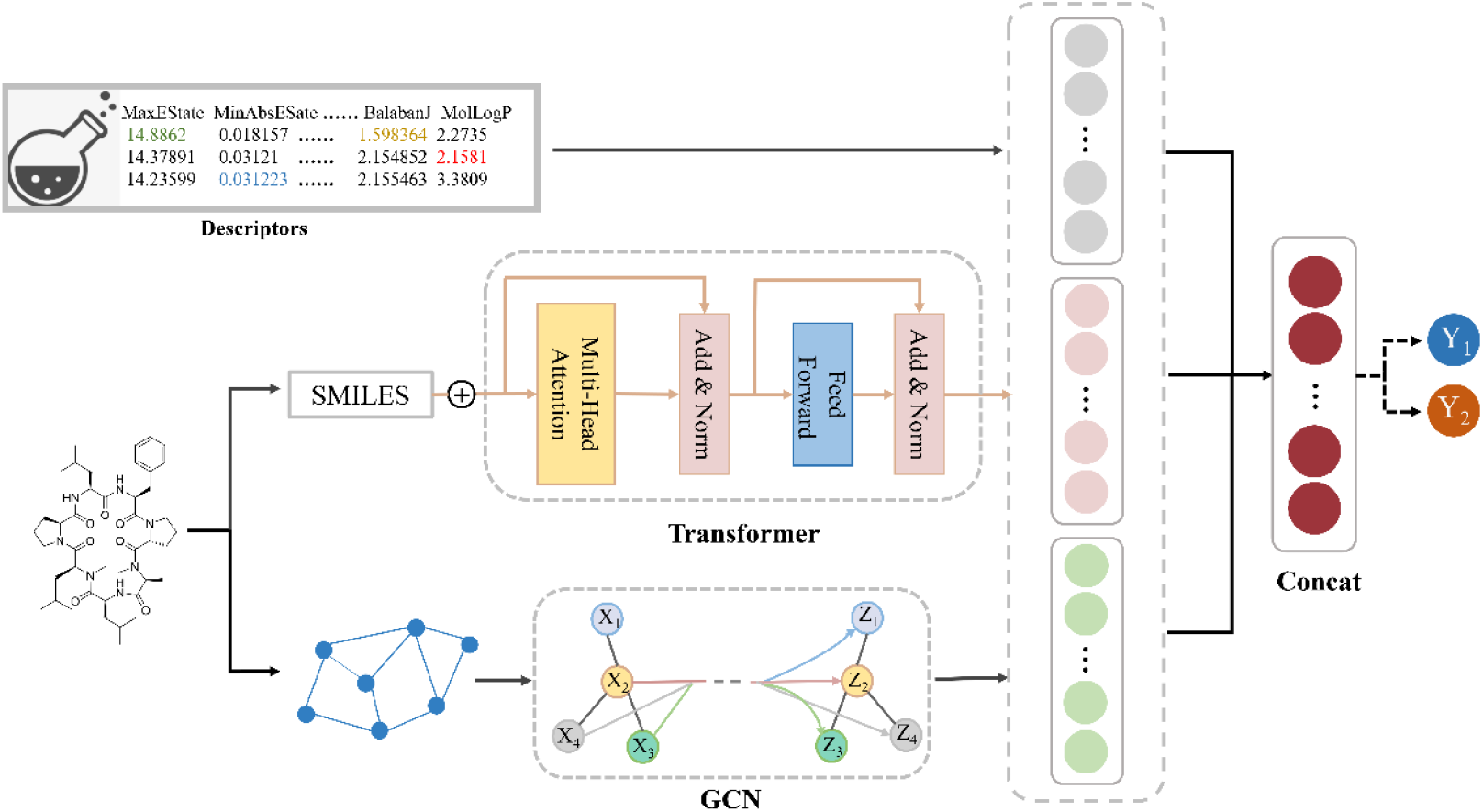
The schematic diagram of the Multi_CycGT model architecture. The main modules of this model are Transformer and GCN.

A specific length of cyclic peptide with a large amount of data can help the model to learn some common feature knowledge, thus improving the model’s performance for other lengths of cyclic peptide in the absence of data. Due to lack of data, the limitations in model performance are not fundamentally addressed, which would result in a large variation in performance across peptide length. We divided the test set into two subsets: length_6/7 and length_other, and results of models with corresponding peptide lengths are shown in Table 2. The prediction indexes for peptides of length_6/7 are generally higher compared to length_other. The multimodal model has an average Recall value of 0.8832 on length_6/7 and 0.7651 on length_other. This indicates that the model performs more accurately in predicting length_6/7, which is consistent with the high proportion of samples of length 6/7 in the training data, meaning that the size of the dataset plays an important role in the training performance of the model. Larger datasets typically provide a larger number of samples and a wider distribution of features, which is a significant advantage to the model.

**Table 2.** The results of Multi_CycGT in 10-fold cross-validation with different metrics.

| Experiment | ACC |  | Pre |  | F1_Score |  | Recall |  | AUC |  |
| --- | --- | --- | --- | --- | --- | --- | --- | --- | --- | --- |
|  | Length | Length_oth | Length_6/ | Length_oth | Length_6/ | Length_oth | Length_6/ | Length_oth | Length_6/ | Length_oth |
|  | _6/7 | er | 7 | er | 7 | er | 7 | er | 7 | er |
| 0 | 0.8359 | 0.6944 | 0.8571 | 0.8920 | 0.8911 | 0.7380 | 0.9280 | 0.6294 | 0.8406 | 0.8397 |
| 1 | 0.8269 | 0.7794 | 0.8750 | 0.8658 | 0.8779 | 0.8255 | 0.8809 | 0.7888 | 0.8548 | 0.8235 |
| 2 | 0.8175 | 0.7847 | 0.8394 | 0.8224 | 0.8630 | 0.8502 | 0.8880 | 0.8800 | 0.8587 | 0.8325 |
| 3 | 0.8046 | 0.7500 | 0.8671 | 0.8688 | 0.8623 | 0.8153 | 0.8576 | 0.7681 | 0.8099 | 0.7986 |
| 4 | 0.7900 | 0.8263 | 0.8321 | 0.9035 | 0.8444 | 0.8768 | 0.8571 | 0.8516 | 0.8353 | 0.8940 |
| 5 | 0.8250 | 0.7426 | 0.9040 | 0.9346 | 0.8750 | 0.8033 | 0.8477 | 0.7044 | 0.8672 | 0.8849 |
| 6 | 0.8293 | 0.7460 | 0.8609 | 0.8918 | 0.8798 | 0.8024 | 0.8996 | 0.7292 | 0.8659 | 0.8380 |
| 7 | 0.8171 | 0.7031 | 0.8196 | 0.9389 | 0.8676 | 0.7639 | 0.9217 | 0.6439 | 0.8445 | 0.8567 |
| 8 | 0.7850 | 0.7830 | 0.8218 | 0.8676 | 0.8401 | 0.8571 | 0.8593 | 0.8468 | 0.8078 | 0.8200 |
| 9 | 0.8200 | 0.7536 | 0.8480 | 0.8222 | 0.8695 | 0.8154 | 0.8921 | 0.8087 | 0.8592 | 0.8225 |
| Average | **0.8151** | **0.7563** | **0.8525** | **0.8808** | **0.8671** | **0.8148** | **0.8832** | **0.7651** | **0.8444** | **0.8410** |

### Conclusion

Peptide membrane permeability prediction is an important and challenging task for the development of cyclic peptide drugs. In this study, we propose a multimodal Multi_CycGT model for cyclic peptide membrane permeability prediction by fusing multi-dimensional features. As the first deep learning-based application for predicting membrane permeability, this method uses GCN and Transformer modules to process features characterizing multiple dimensions (1D and 2D), respectively. Then it splices and fuses them with physicochemical property vector features to finally perform accurate cyclic peptide membrane permeability prediction. We chose to evaluate our model on an independent test set and analytically compare it with currently popular machine learning and deep learning methods such as CNN, Transformer. Extensive cross-sectional experiments demonstrate that Multi_CycGT significantly outperforms those baseline models on four metrics: ACC, Pre, F1, and AUC. This also displays the effectiveness of the multimodal module and highlights Multi_CycGT’s good discriminative ability and generalization capability. In addition, we noticed that there exists the strong implicit relationship between the physicochemical properties of cyclic peptides and membrane permeability. And we hold a point that choosing the correct physicochemical properties is can effectively help the model to understand the potential properties of the peptide. By analyzing the prediction performance of the model with different sequence lengths, we determined that the performance of Multi_CycGT will improve as data increases in the future.

## Method Data

A reliable and rigorous dataset is necessary for the construction and evaluation of statistical prediction models. CycPepMPDB is the first comprehensive transmembrane property database for cyclic peptides. And in the paper, the authors analyze and elaborate on its database^22^. Notably, this database contains three in vitro membrane permeability experiments with more than 200 RDKit generated physicochemical properties. During data preprocessing, we selected 6941 cyclic peptide membrane permeability entries obtained from experiments using the PAMPA method, an in vitro experiment widely used to simulate biological membrane transport. In this experiment, compounds dissolved in the donor pore cross the artificial lipid membrane into the acceptor pore.

The acceptor pore concentration is monitored and the diffusion rate is calculated from the change in acceptor pore concentration over time. In order to normalize the data and improve the performance of models, the physicochemical properties with insufficient information expression, as well as redundant data, were removed. Finally, we obtained a dataset containing 6941 cyclic peptides that have 101 physicochemical properties. Further, we took the cyclic peptides with Permeability>=-6 as the positive samples that could permeate the cell membrane, and the cyclic peptides with Permeability<-6 as the negative samples, and the final ratio of positive to negative samples was 4:6. The data set was randomly divided into training set, validation set and test set by 8:1:1, and an additional 10-fold cross-validation was performed to prevent possible data bias problems to obtain a reliable and stable model.

## Model

### Multi_CycGT architecture

Figure 5 illustrates the detailed framework of the cyclic peptide prediction method Multi_CycGT where the process of predicting cyclic peptide membrane permeability by Multi_CycGT is briefly described. In the multimodal model, considering that the database contains multidimensional features of cyclic peptides, we use different neural network architectures to process the features from different modalities. Specifically, the physicochemical properties were directly turned into vectors by using a fully connected layer, the sequence features were processed by Transformer and the molecular graph features by GCN. After processing the features in three different dimensions, we combine them together to produce a contacted feature vector. Eventually, this integrated feature vector can be fed into our multimodal model for prediction and classification of downstream tasks.

#### Transformer module

The cyclic peptide sequences in the database contain a significant number of non-natural amino acids that undergo modifications based on natural amino acids such as methylation, and amide-ester substitution. By replacing the amino acid sequences with SMILES, the model can acquire information about chemical modifications. Compared to conventional drug small molecules, the SMILES representation of cyclic peptides has several times longer length, which requires the model to handle the longer sequence information properly. Therefore, we utilize the encoder part of the Transformer^24^ to handle the SMILES information of cyclic peptides.

Specifically, each encoder consists of multiple stacked submodules, with each sub-module comprising three components: (1) an embedding layer; (2) a self-attention mechanism layer and (3) a feedforward neural network (FNN). The embedding layer is used to encode the input cyclic peptide SMILES sequence. Given that the position of characters in SMILES is crucial and the molecules characterized by the same token at different positions are completely distinct, a position encoder is employed to incorporate position information of each token into the embedding vector. The position embedding technique used in Transformer combines sine and cosine functions, as follows:

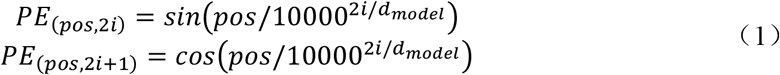

where pos denotes the position, i denotes the embedding vector dimension, and dmodel denotes the dimension of the model. The sine and cosine functions with different frequencies are used to encode each position, thus enabling the model to capture information about the relative distances between different positions in the input sequence.

The attention mechanism is the core of the whole encoder. Each token in the sequence will be match with features of other tokens as keys based on its own features as a query, and finally a weighted representation of the context is obtained. It can pay attention to the important information in a large amount of information and reduce the influence of redundant information. And the self-attention mechanism, as a variation of the attention mechanism, captures the internal association of sequences and reduces the dependence on external information. Its calculation process is shown as follows:

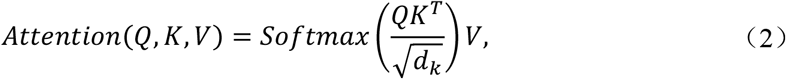

where the three matrices Q(Query), K(Key) and V(Value) are generated based on the input sequence.

The calculation process can be summarized as three processes: (1) First, the dot product between Q and K is calculated, with a subsequent division by the square root of the dimension of the Key vector (denoted as dk) to prevent excessively large results. (2) The resulting values are normalized using the Softmax operation, generating a probability distribution. This distribution is then multiplied by the matrix V to obtain a representation of the weight summation. The attention score indicates the importance of the information, with higher scores signifying greater focus on relevant details. Through the utilization of a multi-layered self-attention mechanism, the encoder can learn different representation subspaces, facilitating the extraction of more comprehensive semantic information from the input sequence.

The feedforward neural network consists of multiple fully connected layers and incorporates residual connections to refine feature representation and predict the final permeability score. This network architecture aids in capturing intricate relationships and enhancing the overall modeling capacity of the system.

#### Graph Convolutional Networks (GCN)

A graph convolutional network (GCN) is a neural network specifically designed to process data with graph structure, such as molecular graphs. It is particularly useful for analyzing the spatial relationships among atoms or substructures within cyclic peptides.

In the graph representation of cyclic peptide molecules, the nodes represent the atoms in the graph, while the edges represent the chemical bonds connecting the atoms. In GCNs, the atom connections are treated as adjacency matrices, and each node in the graph is initialized with a feature vector that encodes relevant atom-features characteristics, including atom_type_one_hot, atom_degree_one_hot, atom_implicit_valence_one_hot, atom_ formal_charge, atom_num_radical_electrons, atom_hybridization_one_hot, atom_is_aromatic, and atom_total_num_H_one_hot. The central operation in GCNs is the message passing step, which is usually performed through several graph convolution layers. Each layer updates node features based on the aggregated information of neighboring nodes. These layers capture increasingly complex relationships and dependencies among atoms within the peptide. The computational equation for GCNs can be expressed as follows:

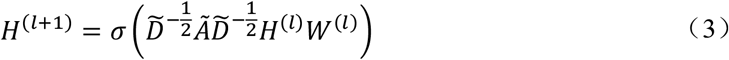

where *H*^*l*^ is the layer l network output, *Ã* is the adjacency matrix, 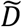 is the degree matrix of the adjacency matrix, *W*^(*l*)^ is the trainable weight matrix, and *σ* is the activation function.

The information at the (l+1)th layer of each node is obtained by weighted summation of the information at the lth layer with the information of neighboring nodes, which is then transformed linearly and nonlinearly.

By employing the GCN, the two-dimensional structural features present in the molecular graph can be effectively captured and processed. The inherent ability of GCNs to incorporate graph structure information allows for a comprehensive analysis of the relationships and patterns among atoms within the cyclic peptide. This enables the extraction of meaningful features that can be utilized for subsequent classification tasks^25^.

### Baselines

Except for Transformer and GCN, we compare our model with four different models in this article.

#### Support Vector Machines (SVM)

SVM^26^ is a powerful and widely used machine learning model, and it is commonly applied to both classification and regression problems. In the context of classification, SVM aims to find an optimal hyperplane that maximally separates different classes in the feature space. It achieves this by identifying a subset of training samples, known as support vectors, that are closest to the decision boundary.

#### K-Nearest Neighbors (KNN)

KNN^27^ is a versatile algorithm commonly used for data mining and image classification problems. It works by classifying data points based on the majority vote of their nearest neighbors in the feature space.

#### Convolutional Neural Network (CNN)

CNN^28, 29^ is a type of feedforward neural network that incorporates convolutional operations and typically has a deep architecture. CNNs are widely recognized as one of the prominent algorithms in the field of deep learning. The convolutional layers in a CNN perform feature extraction through the application of filters or kernels, followed by pooling layers for down-sampling. The deep structure of CNNs enables them to learn hierarchical representations of the input data, making them highly effective in various tasks.

#### Long Short-Term Memory (LSTM)

LSTM^30^ networks are a specific type of Recurrent Neural Network (RNN) architecture that excels in capturing and learning long-term dependencies in sequential data. Unlike traditional RNNs, LSTM networks are designed to mitigate the vanishing gradient problem, which can hinder the ability of RNNs to retain information over long sequences. LSTMs achieve this by utilizing memory cells and gates that regulate the flow of information, allowing them to selectively remember or forget information at different time steps.

### Evaluation Metrics

In order to avoid overfitting or underfitting, 10-fold cross-validation is chosen in this paper to assess the prediction performance of the method. Meanwhile, some effective evaluation metrics are selected to verify the feasibility of the model. They are accuracy (ACC), precision (Pre), F1-score (F1), recall score (Recall) and area under the curve (AUC), respectively. The correlation equation for the four metrics is expressed as follows:

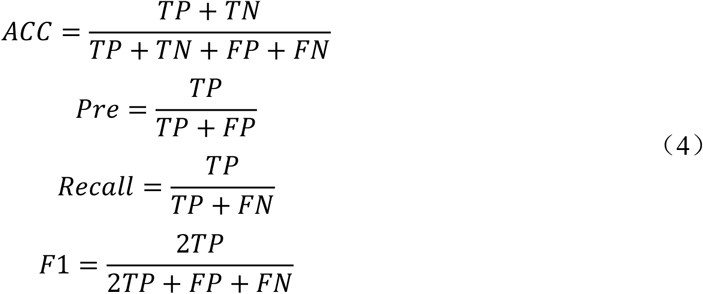

where TP (true positive), TN (true negative), FP (false positive), and FN (false negative) are used to represent the respective counts of each category. Additionally, ROC (Receiver Operating Characteristic) curves and AUC are widely employed for assessing performance. The metrics have values ranging from 0 to 1, and higher values of these metrics indicate better performance of the model.

## Funding Information

This project was supported by the Natural Science Foundation of Zhejiang Province (LD22H300004).

## Notes

### Competing Interest Statement

The authors have declared no competing interest.

